# Platelet-specific TGFβ1 deficiency aggravates atherosclerosis, vascular inflammation, and hypercholesterolemia in mice

**DOI:** 10.1101/2023.10.20.563268

**Authors:** Shuai Tan, Yang Sun, Zi Sheng, Yanan Min, Anton Gisterå, Junhao Zhang, Daniel F.J. Ketelhuth, Wangjun Liao, John Andersson, Hu Hu, Miao Wang, Ming Hou, Mingxiang Zhang, Jun Peng, Chunhong Ma, Nailin Li

## Abstract

Atherosclerosis involves inflammatory and thrombotic mechanisms, to which both platelets and transforming growth factor β (TGFβ) contribute. The effect of platelet-derived TGFβ on atherosclerosis is, however, unknown and therefore investigated. Murine platelet-selective TGFβ-deficiency (plt-TGFβ^-/-^) was created by a *Pf4*-Cre approach, and an atherosclerotic mouse model was established by functional abrogation of *Ldlr* and 10-15 weeks of a high-fat diet in plt-TGFβ^-/-^ mice and their non-plt-TGFβ^-/-^ littermates. En face Oil Red O staining of the aorta showed more atherosclerotic lesion formation in plt-TGFβ^-/-^ mice, with significant increases in both lesion size and lesion coverage of the total aortic area. Cryosections of the aortic root confirmed the aggravation of atherogenesis. Platelet-derived TGFβ deficiency increased circulating platelets and plasma levels of total cholesterol, LDL-cholesterol, and triglycerides after a 10 or 15 week high-fat diet period. RNA sequencing and proteomic analyses of the aorta showed signs of CD4^+^ T effector cell and macrophage activation in plt-TGFβ^-/-^ mice. In conclusion, platelet-specific TGFβ deficiency aggravates atherosclerosis, via increasing arterial inflammation and plasma levels of cholesterol. Our findings demonstrate that platelet-derived TGFβ is prominently athero-protective.

**Key points:**

- Platelet-specific transforming growth factor β (TGFβ) deficiency markedly enhances atherosclerosis in a high-fat diet-fed murine model.
- Platelet TGFβ deficiency aggravates hyperlipidemia, with further elevations of total cholesterol, LDL-cholesterol, and triglycerides.

## Introduction

Platelets and CD4^+^ T cells (e.g., type 1 T helper/Th1 and Th17 cells and regulatory T/Treg cells) are closedly involved in atherosclerosis.^1-5^ Transforming growth factor β (TGFβ) plays an important protective role in atherosclerosis.^3-5^ Platelets are the principal source of TGFβ in circulation,^6^ and can regulate various aspects of CD4^+^ T cell activities.^4,7^ The impact of platelet-derived TGFβ on atherogenesis is, however, unknown. We hypothesized that platelet-specific TGFβ deficiency (plt-TGFβ^−/−^) would aggravate atherogenesis.

## Methods and Materials

### Ethical consideration

The study was performed with a pro-atherosclerotic murine model combining plt-TGFβ^−/−^, high-fat diet, and LDLR deficiency. All relevant ethical guidelines and animal welfare regulations for animal studies have been followed. All necessary ethics committee approvals have been obtained (see below).

### Mice

All animal study protocols have been reviewed and approved by the Animal Ethics Committee of Shandong University Cheeloo Medical College (Jinan, China; Dnr SDUQH 2016-10-13-1). All mice were bred and housed at the animal research center of the Cheeloo Medical College in specific pathogen-free conditions.

All genetically modified mice were created on a C57BL/6 background. Platelet-specific TGFβ1 deficient (Plt-TGFβ^−/−^) mice were established by crossing mice carrying a “floxed” TGFβ1 allele (Tgfb1^flox^; Jackson Laboratory) to mice expressing Cre recombinase under the control of the megakaryocyte/platelet-specific *Pf4* promoter (*Pf4*-Cre^+^ transgenic mice) (Model Animal Research Center of Nanjing University, Nanjing, China). Eight-week-old male mice were used for the experiments.

### Murine model of atherosclerosis

Hypercholesterolemic mice were created by a gain-of-function approach by a tail vein injection of proprotein convertase subtilisin/kexin type 9 (PCSK9)-encoding rAAV8 vectors^8^ (1.0×10^11^ vector genomes/mouse) in 8-week-old plt-TGFβ^−/−^ mice (Pf4-Cre^+^-TGF-β1^f/f^) and their littermate controls (Pf4-Cre^-^-TGF-β1^f/f^). After 1-week-incubation allowing the functional abrogation of LDLR, the mice were fed with a cholate-free Western-type high-fat diet (21% fat and 0.21% cholesterol) for 10 to 15 weeks. The mice were then sacrificed. Blood was collected by cardiac puncture with a 1 ml EDTA-vacutainer. The right auricle was cut, and 5 ml cold phosphate-buffered saline (PBS) was slowly infused via the left ventricle. The heart, the entire aortic tree, the spleen, and the lymph nodes were then harvested under an anatomy microscope.

### Blood analyses

Blood samples were collected from the tail vein at baseline. Murine EDTA whole blood samples were used for blood cell counting (Mindray; Shenzhen, China). Plasma was separated by centrifugation at 1,500*g* for 15 min at 4°C. The plasma samples were aliquoted at 60 µl/tube and stored at -80°C for subsequent analysis. Lipids were assayed in plasma at the basal line (8-week from birth) and after 10-week or 15-week on a high-fat diet. The plasma level of TGFβ1 was measured using a mouse TGFβ1 precoated ELISA kit (Cat: 1217102; ThermoFisher) and a Tecan microplate reader (Infinite M Nano; Männedorf, Switzerland).

### Flow cytometry

The spleen and lymph nodes were mashed using slides. The cells were resuspended with cold PBS and filtered. Splenocytes underwent red cell lysis using BD PharmLyse™ Lysing Buffer. The cells were then washed with PBS and spun at 400*g*. The cells were then resuspended in PBS at the concentration of 10^7^ cells/ml. For cell surface marker staining, 100 µl cell suspension (10^6^ cells) was stained with corresponding fluorescent antibodies for 30 min at 4°C. For intracellular staining, the cells were seed on 12-well plates and stimulated with PMA (50 ng/ml) and ionomycin (5 µg/ml) for 2 hours, followed by incubating with Brefeldin A for 4 hours to block cytokines secretion. Afterward, the cells were fixed with a Fixation Buffer (Cat#00-8222-49; ThermoFisher, USA) and permeabilized using a Permeabilization Buffer (Cat#00-8333-56; ThermoFisher), and then washed and stained with corresponding fluorescent antibodies for 30 min at 4°C. The stained cells were resuspended in PBS and analyzed using CytoFLEX S or Galios® flow cytometers (Beckman-Coulter Corp., Hialeah, FL, USA). Data analyses were performed using FlowJo version 10.

The antibodies used in the staining were PECy7-CD3 (clone 17A2, Cat#100220; Biolegend, San Diego, CA, USA), PerCPCy5.5-CD4 (clone RM4-5, Cat#45-0042-82; eBioscience, San Diego, CA, USA), BV421-IFN-γ (Cat#11-9668-82; eBioscience), PE-IL-4 (clone 11B11, Cat#554435; BD Pharmingen, San Jose, CA, USA), FITC-IL17A (clone BL168, Cat#512304; Biolegend), APC-T-bet (Cat#130-098-655; Miltenyi Biotec, Bergisch Gladbach, Germany), FITC/Alexa488-CD25(clone 3C7, Cat#53-0253-82; eBioscience), PECy7-CD39 (clone DuHa59, Cat#143806; Biolegend), eFlour660-FoxP3 (clone FJK-16s, Cat#50-5773-82; eBioscience), PE-TGF-beta1 (latency-associated peptide/LAP) (clone TW7-20B9, Cat#141305; Biolegend), PE-CD278 (ICOS) (clone C398.4A, Cat#313508; Biolegend), FITC-CD185 (CXCR5) (clone L138D7, Cat#145520; Biolegend), APCCy7-CD279 (PD-1) (clone 29F.1A12, Cat#135224, Biolegend).

### Cryosectioning, Oil red O (ORO) staining, and en face analysis

The aortic roots in OCT were mounted on the specimen holder in a cryostat-microtome, and the proximal 800 μm of the aortic root was serially sectioned as described previously.^9^ Cryosections (10 µm) were collected on Superfrost Plus microscope slides (Thermo Scientific, cat. no. J1800AMNZ). Cryosections were fixed in cold 100% acetone for 10 min, air-dried at room temperature for 30 min, and then stored at -20°C until staining.

Lipid staining was performed using an ORO staining solution (5 mg/ml ORO in isopropanol, and 3:2 diluted with water before use). The dissected thoracic aorta samples were inserted into 2 ml ORO solution and stained for 3 hours at room temperature. The samples were then washed with 70% ethanol and stored in PBS at 4°C. After pinning on a wax plate, the aortas were photographed as previously described.^10^ The cryosections of the aortic root were stained in ORO solution for 20 min at room temperature. Afterward, the sections were rinsed with water, re-stained with Hematoxylin QS for 40 s, rinsed again with water, and mounted with a water-based mounting medium. Micrographs of the cryosections were obtained using an OLYMPUS BX61VS microscope. Lesion size/area of the aorta and the cryosections were analyzed using the programs OlyVIA Version 2.9 and ImageJ 64bit.

### RNA sequencing and proteomic analyses

Total RNA of a separate batch of aortas, not used for en face lesion quantification, was extracted using RNeasy Mini Kit (Cat#74104; Qiagen, Hilden, Germany) according to the manufacturer’s instruction. After taking the RNA-containing upper fraction of the Qiazol reagent, the pink organic fraction was used for protein isolation. Briefly, absolute ethanol was added to the pink organic fraction, thoroughly mixed, and incubated at room temperature for 5 min. DNA was then sedimented by centrifugation (2,000 *g*, 5 min, 4°C). The protein-containing supernatant was collected, and proteins were precipitated by adding isopropanol and incubated at room temperature for 10 min. After centrifugation (12,000 *g*, 10 min, 4°C) and removing the supernatant, the protein pellet was washed with 0.3 M Guanine HCL in 95% ethanol, and then stored at -80°C.

The RNA sequencing and mass spectrometry/proteomics analyses were performed by the Beijing Genomics Institute (BGI). The data were analyzed using the Dr. Tom program. The RNA sequencing data have been deposited to NCBI bioproject site with the accession number PRJNA990504. The mass spectrometry proteomics data have been deposited to the ProteomeXchange Consortium via the PRIDE [1] partner repository with the dataset identifier PXD043833 (Username; Password: YrfNNLf7).

### Statistical analysis

The data were are presented as mean±SEM. Student’s *t-*test and Wilcoxon signed-rank test were used to compare to compare the differences between groupstwo treatments. Multiple comparisons between the treatments were analysed analyzed by one-way ANOVA followed by post hoc Tukey tests using SPSS 22 and Graphpad Prism 5. A *p*-value!0.05 was considered statistically significant.

## Results and Discussion

The present study found that plt-TGFβ^-/-^ mice display markedly aggravated atherosclerosis, with enhanced lesion formation, altered CD4^+^ T effector activities in peripheral lymphoid organs, increased circulating platelets, as well as elevated plasma lipid levels.

### Plt-TGFβ^-/-^ aggravates atherosclerosis

En face ORO staining of the aorta demonstrated that plt-TGFβ^-/-^ markedly increased atherosclerotic lesion formation after 10-week high-fat diet, as compared to non-TGFβ1 deficient littermates, and that the increase was more prominent in the aortic arch and the thoracic aorta and even more obvious after 15 weeks (figure 1A). The aggravated lesion formation was seen both in lesion size and lesion coverage (approximately from 2 to 5%; figure 1B). Smaller lesions were formed in the abdominal aorta, and atherosclerotic lesion area tended to higher in plt-TGFβ^-/-^ mice than in control mice (0.59±0.14mm^2^ *vs*. 0.24±0.07mm^2^, *p*=0.053). ORO staining of the aortic root cryosections showed significant atherosclerotic lesion formation in plt-TGFβ^-/-^ mice (figure 1C), both in terms of absolute lesion area and relative lesion area (figure 1D).

**Figure 1.**
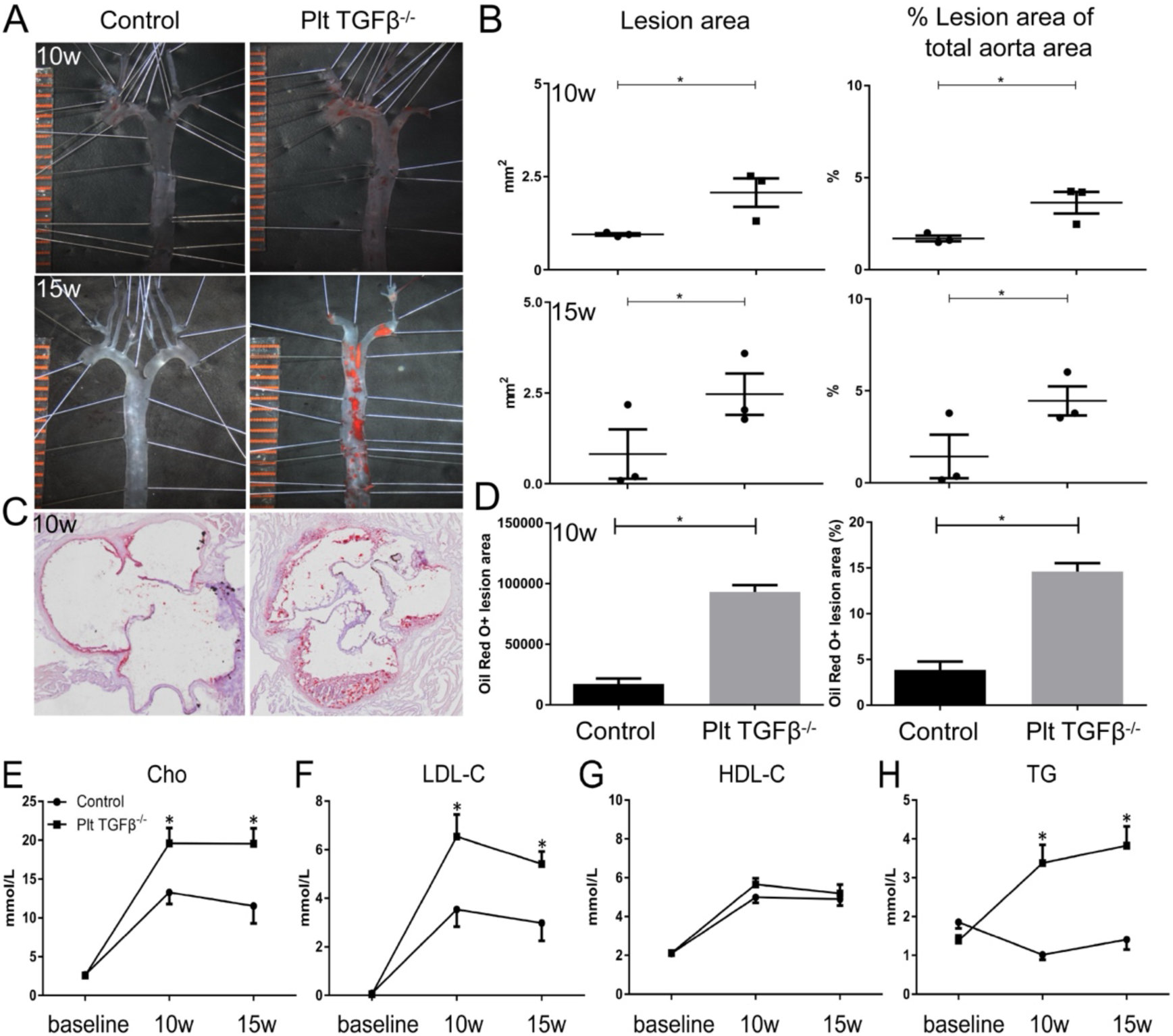
Platelet-specific TGFβ deficiency exaggerates hyperlipidemia and aggravates atherosclerosis. Platelet-specific TGFβ1 deficient (Plt-TGFβ^−/−^) mice were established by crossing mice carrying a “floxed” TGFβ1 allele (*Tgfb1*^flox/flox^) to mice expressing Cre recombinase under control of the megakaryocyte/platelelet-specific Pf4 promoter (Pf4-Cre^+^). The pro-atherosclerotic mice of LDLR-functional knockout were created using a gain-of-function approach by overexpression of proprotein convertase subtilisin/kexin type 9 (PCSK9) in 8-week-old plt-TGFβ^−/−^ male mice (*Pf4*-Cre^+^-*Tgfb1*^f/f^) and their littermate control (*Pf4*-Cre^-^-*Tgfb1*^f/f^). The mice were feed with a cholate-free Western-type high-fat diet (21% fat and 0.21% cholesterol) for 10 to 15 weeks. After the mice were sacrificed and the cardiovasculature was perfused with cold PBS, the heart and the aorta were harvested. Panel A: En face staining of atherosclerotic lesions. The aortas were fixed with 4% paraformaldehyde in PBS at 4°C for ≥24 h, stained with 3 mg/ml ORO at room temperature for 3 h, washed with 70% ethanol, and stored in PBS at 4°C. The aortas were then pinned open on a black surface and photographed. Representative images of the aortas from the mice after 10-week (upper images) and 15-week high-fat diet were shown. Scale bar showed orange mm-marks. Panel B: Quantification of atheroclerotic burden in the en face prepared aortas. The images of ORO-stained atherosclerotic lesions were analysed using Image J programme. Lesion area in mm^2^ and the percentages of atherosclerotic lesions in the total aorta face area are presented. * *p*<0.05, as compared to the control mice; n=3. Panels C-D: Atherosclerotic lesions of the aorta root. ORO-stained atherosclerotic lesions in cryosections from the aortic root, 300 µm from the aortic sinus, in mice fed a high-fat diet for 10 weeks. The cryosections were observed and photographed using an OLYMPUS BX61VS microscope. Lesion area is presented in arbitrary units and as a percentage of total vessel circumference. * *p*<0.05, plt-TGFβ^−/−^ vs. control mice; n=3-6. Panels E-H: Plasma samples at baseline and after 10 w and 15 w on a high-fat diet were collected for analyses of the total cholesterol (Cho; panel E), low-density lipoprotein-cholesterol (LDL-C; panel F), high-density lipoprotein-cholesterol (HDL-C; panel G), and triglycerides (TG; panel H) (One-way ANOVA followed by post hoc Tukey test, \**p*<0.05, n=3-9).

### Plt-TGFβ^-/-^ elevates lipid levels and platelet counts

Plasma levels of TGFβ were β70% and persistently reduced in plt-TGFβ^-/-^ mice (Supplementary figure 1), indicating the successful knockout of platelet-derived TGFβ and platelets as the principal source of blood TGFβ.

Functional LDLR-knockout by PCSK9 overexpression and high-fat-diet feeding induced hypercholesterolemia in the mice, with elevated total cholesterol (Cho), low-density lipoprotein cholesterol (LDL-C), but not triglycerides (TG) (figure 1E-H). Interestingly, plt-TGFβ^-/-^ further elevated the plasma levels of Cho and LDL-C, but not HDL-C. On the other hand, plt-TGFβ^-/-^ also increased TG levels, which were not increased by LDLR deficiency or high-fat diet. Together, these results highlighted that plt-TGFβ^-/-^ elevated LDL-C and TG levels, which commonly contribute to increased atherosclerotic lesion formation,^5,11^ and that lipid-lowering mechanisms by platelet-dervied TGFβ may anchor to the key lipid metabolic organ liver and require further delineation.

Plt-TGFβ^-/-^ mice had elevated platelet counts, which may reflect enhanced thrombotic and inflammatory states. Plt-TGFβ^-/-^ increased platelet counts from 669±115×10^9^/L in control mice to 885±52×10^9^/L (*p*<0.05). The latter was further elevated by the high-fat diet, to 1006±112×10^9^/L at 10 weeks and to 1222±74×10^9^/L at 15 weeks (*p*<0.05, n=5-9). These data suggest that platelet-derived TGFβ may serve as a feedback regulator of platelet generation, and confirm the correlation between elevated PCSK9/LDL levels and platelet count/hyperreactivity.^12^

### Plt-TGFβ^-/-^ alters CD4^+^ effector cell activities in the spleen and lymph nodes

Platelet-derived TGFβ is the major source of circulating TGFβ,^6^ and is an important regulator of CD4^+^ T effector responses.^4^ CD4^+^ T effector cell phenotyping (supplementary figure 2A) showed that plt-TGFβ^-/-^, unexpectedly, decreased the percentages of Th1 and Treg cells (figure 2A-D), as well as Th2 and Th17 cells (supplementary figure 2B-E) in both the spleen and the lymph nodes in the 10-week high-fat diet-fed mice. Later during the disease, however, Th1/T-bet expression significantly increased in lymph node cells (figure 2F) and tended to increase in splenic CD4^+^ T cells (*p*=0,10; figure 2E), which was consistent with the Th1 phenotype/IFNγ content (figure 2G&H). In contrast, while Treg/FoxP3 expression was similar between plt-TGFβ^-/-^ and control mice (figure 2I&J), intracellular levels of TGFβ, a marker of Treg cell immune-suppressing activities, were markedly reduced in plt-TGFβ^-/-^ mice (figure 2K&L). Expression of another Treg cell anti-inflammatory marker CD39, an ectonucleotidase that cleaves adenosine triphosphate (ATP), on Treg cells was also decreased in plt-TGFβ^-/-^ mice (figure 2M&N). Our data suggest that effector T cells may secrete increased amounts of the proatherogenic cytokine IFNγ, and that Treg cell functionality might be compromised in plt-TGFβ^-/-^ mice.

**Figure 2.**
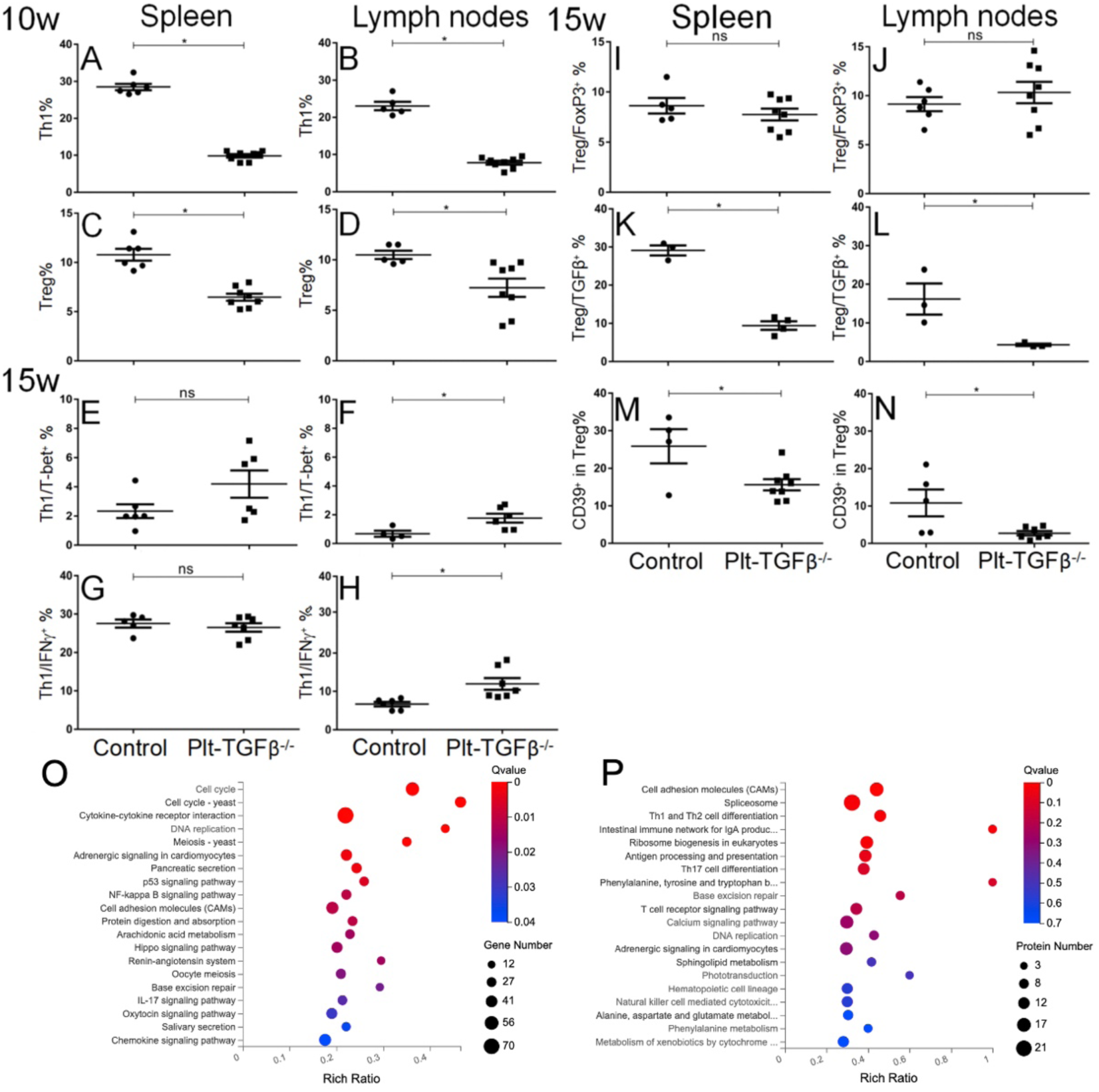
Platelet-specific TGFβ deficiency alters inflammatory landscape in the aorta and CD4^+^ effector cell status in the spleen and lymph nodes. Plt-TGFβ^−/−^ mice and their littermate control were fed with a high-fat chow for 10 and 15 weeks. The spleen and lymph nodes were harvested for the preparation of single cell suspensions at both time points. Flow cytometric phenotyping of CD4^+^ T effector cells, which were first stimulated using PMA (50 ng/ml) and ionomycin (5 µg/ml) for 2 hours, and followed by adding brefeldin A (5 µM) for further incubation for 4 hours. Student’s *t*-test, \**p*<0.05, n=3-9. Panels A-D: CD4^+^ T effector cell phenotypes after 10-w high-fat diet. Th1 cells gated as CD4^+^-IFNγ^+^ cells, and Treg cells as CD4^+^-CD25^++^-FoxP3^+^ from the spleen (A and C) and the lymph nodes (B and D). Panels E-H: Th1 cells phenotyped by T-bet and by IFNγ staining from the cells of the spleen (E and G) and the lymph nodes (F and H) after 15-w high-fat diet. Panels I-N: Treg cells were phenotyped by CD25-FoxP3, TGFβ, and CD39 staining from the cells of the spleen (I" K" and M) and the lymph nodes (J" L and N) after 15-w high-fat diet. Mean±SEM were plotted, \**P*<0.05 compared to control by Student’s *t*-test, n=4-8. Panels M and N: Plt-TGFβ^−/−^ mice and littermate controls were fed a high-fat diet for 15 weeks. The whole aorta was harvested, snap-frozen, and stored at -80°C. The Qiazol reagents were used for the total RNA and protein isolation. The aqueous phase was used for RNA isolation, while DNA was sedimented by pure ethanol. Protein isolation was performed by isopropanol precipitation of the organic phase. RNA sequencing was performed using the DNBSEQ™ Technology Platform, and mass spectrometric analyses and proteome profiling were carried out at BGI (Shenzheng, China). Data were analysed with the Dr. Tom program. Kyoto encyclopedia of genes and genomes (KEGG) pathway analysis of RNA sequencing (O) and mass spectrometrics analysis (P) were presented.

### Plt-TGFβ^-/-^ elicits a pro-inflammatory profile in the aortic vessel wall

To further reveal plt-TGFβ^-/-^ impact on vascular inflammation, RNA sequencing data from the aortas of mice on 15-week high-fat diet demonstrated that plt-TGFβ^-/-^ promoted the activation of multiple inflammatory pathways, such as cytokine-cytokine receptor interactions, IL-17 signaling pathway, chemokine signaling pathways, and arachidonic acid metabolism (Kyoto encyclopedia of genes and genomes/KEGG pathway analyses, figure 2O). Gene ontology (GO) enrichment analyses also showed that plt-TGFβ^-/-^ enhanced lymphocyte and monocyte chemotaxis/infiltration and inflammatory responses (supplementary figure 3A).

Inline with the above findings, proteomic data demonstrated that plt-TGFβ^-/-^ promoted antigen processing and presentation and enhanced CD4^+^ T effector cell responses (figure 2P), and increased the expression of the major histocompatibility complex class II (MHC-II) protein complex (supplementary figure 3B). Together, transcriptomic and proteomic analyses supported that plt-TGFβ^-/-^ enhanced inflammatory activities in the vessel, especially monocyte/macrophage-dependent antigen processing/presentation and CD4^+^ T cell-driven effector cell responses.

There are several interesting findings here. Firstly, we show that platelet-derived TGFβ is critical in controlling vascular inflammation and is athero-protective. Specifically, plt-TGFβ^-/-^ resulted in a clear pro-inflammatory landscape in the aorta, with coherent findings from the transcriptomic/proteomic analyses and aggravated atherosclerosis. TGFβ is largely considered athero-protective.^13^ Platelets selectively adhere on and infiltrate into the atherosclerotic lesion,^4,14^ and secrete TGFβ to skew T cell responses to enhance anti-inflammatory Treg cell activities, but inhibit pro-inflammatory Th1 effector responses.^4^ Platelet-derived TGFβ likely influences other vascular cells, such as smooth muscle cells and monocytes/macrophages, towards stabler plaques.^13^ These diverse TGFβ actions collectively transform into an athero-protective effect, although not all of them were explored in the present study. Notably, TGFβ may evoke pro-atheroclerotic vascular inflammatory responses of endothelial cells, and endothelial TGFβ signalling blockade may reduce vascular inflammation and atherosclerosis,^15,16^ highlighting the cell-type-specific impact of TGFβ on atherogenesis.

Another interesting finding was that plt-TGFβ^-/-^ significantly elevated plasma levels of lipids. LDLR deficiency and a high-fat diet elevated, as expected, the levels of total cholesterol, HDL-C, and LDL-C. Remarkably, plt-TGFβ^-/-^ further enhanced the levels of total cholesterol and LDL-C, but not HDL-C. We have previously shown that platelet-derived TGFβ enhances Treg cell proliferation and responses,^4,17^ and that Treg cells modulate blood lipid levels by enhancing lipoprotein catabolism in another hypercholesterolemic mouse model.^18^ Hence, decreased Treg cell differentiation and especially hampered Treg cell functions by plt-TGFβ^-/-^ may lead to hyperlipidemia. Furthermore, platelets closely regulated hepatocyte and liver functions.^19^ TGFβ triggers multiple signaling, e.g., phosphorylated-Smad2/3-dependent and mitogen-activated-protein kinase, pathways in hepatocytes^20^ that can regulate the activation of the transcription factor SREBP1/2 and thus control not only PCSK9 but also LDLR synthesis.^21,22^ Considering the marked elevation of lipid levels by plt-TGFβ^-/-^, we propose that platelet-derived TGFβ may enhance LDLR synthesis, attenuate PCSK9 production, and subsequently suppress lipid levels in circulation.

The present study had limitations. The processing of aortas for transcriptomic and proteomic analysis provided no materials for the investigation of inflammatory cells in situ by immunohistochemical staining. This was not possible in aortic root sections either, due to technical difficulties.

In conclusion, platelet-specific TGFβ deficiency aggravates atherosclerosis by augmenting vascular inflammation and hyperlipidemia. The findings highlight that therapeutic approaches to potentiate platelet-derived TGFβ activities could be a promising strategy for atherothrombotic disease management.

## Acknowledgements

The study was supported by grants from the Swedish Heart-Lung Foundation, the National Natural Science Foundation of China (project no 81700110), the Shandong University-Karolinska Institutet Cooperative Research Fund, the Swedish Foundation for Internationalisation of Higher Education and Research (STINT), the China Scholarship Council, Karolinska Institutet, and the Stockholm County Council.

## Author contributions

S. Tan, Y. Sun, and Z. Sheng performed experiments, interpreted the data, and wrote the manuscript; Y. Min performed research, interpreted the data, and revised the manuscript; A. Gisterå and J. Zhang performed experiments, interpreted the data, and revised the manuscript. D.F.J. Ketelhuth, W. Liao, J. Andersson, H. Hu, M. Wang, M. Hou, M. Zhang, and J. Peng designed the study, interpreted the data, and revised the manuscript; C. Ma and N. Li designed the study, interpreted the data, organized the research, and wrote the manuscript.

