## Supplementary materials for "Platelet-specific TGFβ1 deficiency aggravates atherosclerosis, vascular inflammation, and hypercholesterolemia in mice"

[1]Karolinska Institutet, Department of Medicine-Solna, Cardiovascular Medicine Unit, Stockholm, Sweden; [2]Shandong University Cheeloo Medical College, School of Basic Medicine, Department of Immunology, Jinan, China; [3]Shandong University-Karolinska Institutet Translational Medicine Collaborative Research Platform, Jinan, China; [4]Qilu Hospital of Shandong University, Department of Hematology, Jinan, China; [5]Jining Medical University Affiliated Hospital, Department of Hematology, Jining, China; Karolinska Institutet, [6]Center for Molecular Medicine, Karolinska University Hospital, Stockholm, Sweden; [7]Nanfang Hospital, Department of Oncology, Southern Medical University, Guangzhou, China; [8]Department of Cardiovascular and Renal Research, Institute for Molecular Medicine, University of Southern Denmark; [9]Department of Environmental Medicine, Integrated Epidemiology Group, Stockholm, Sweden; [10]Zhejiang University School of Medicine, Department of Pathology and Pathophysiology, Hangzhou, China; [11]Fuwai Hospital, National Center for Cardiovascular Diseases, Chinese Academy of Medical Sciences and Peking Union Medical College, Beijing, China; [12]Qilu Hospital of Shandong University, Key Laboratory of Cardiovascular Remodeling and Function Research, Jinan, China

### Abstract

Atherosclerosis involves inflammatory and thrombotic mechanisms, to which both platelets and transforming growth factor β (TGFβ) contribute. The effect of platelet-derived TGFβ on atherosclerosis is, however, unknown and therefore investigated. Murine platelet-selective TGFβ-deficiency (plt-TGFβ$^{-/-}$) was created by a *Pf4*-Cre approach, and an atherosclerotic mouse model was established by functional abrogation of *Ldlr* and 10-15 weeks of a high-fat diet in plt-TGFβ$^{-/-}$ mice and their non-plt-TGFβ$^{-/-}$ littermates. En face Oil Red O staining of the aorta showed more atherosclerotic lesion formation in plt-TGFβ$^{-/-}$ mice, with significant increases in both lesion size and lesion coverage of the total aortic area. Cryosections of the aortic root confirmed the aggravation of atherogenesis. Platelet-derived TGFβ deficiency increased circulating platelets and plasma levels of total cholesterol, LDL-cholesterol, and triglycerides after a 10 or 15 week high-fat diet period. RNA sequencing and proteomic analyses of the aorta showed signs of CD4$^{+}$ T effector cell and macrophage activation in plt-TGFβ$^{-/-}$ mice. In conclusion, platelet-specific TGFβ deficiency aggravates atherosclerosis, via increasing arterial inflammation and plasma levels of cholesterol. Our findings demonstrate that platelet-derived TGFβ is prominently athero-protective.



**Keywords**: Platelets; Transforming Growth Factor β, CD4$^{+}$ T cells, Atherosclerosis, Cholesterol

### Supplementary Results

#### ***Platelet-specific TGFβ1 deficiency reduces circulating TGFβ levels***

Platelet-targeted knockout of TGFβ resulted in significant reduction of circulating TGFβ in plt-TGFβ$^{-/-}$ mice as compared to their control littermates, and similar reductions were seen both after 10-W and 15-W high-fat diet (Supplementary figure 1).

#### ***Plt-TGFβ$^{-/-}$ alters CD4$^+$ effector cell activities in the spleen and lymph nodes***

Flow cytometric phenotyping of CD4$^+$ effector cells were performed using intracellular staining, in which IFNγ$^+$, IL-4$^+$, IL-17A$^+$, and FoxP3$^+$-CD25$^{++}$ were used as the markers of Th1, Th2, Th17, and Treg cells among CD4$^+$ cells, respectively (Supplementary figure 2A). It was found that plt-TGFβ$^{-/-}$ decreased the percentages of Th2 (Supplementary figure 2B&C) and Th17 (Supplementary figure 2D&E) among the CD4$^+$ T cells from both the spleen and the lymph nodes in the 10-week high-fat diet-fed mice. When the duration on high-fat diet was extended to 15 weeks, the suppressed CD4$^+$ effector cells were seen in Th2 cells from the lymph nodes (Supplementary figure 2H) and Th17 cells in the spleen (Supplementary figure 2I).

With the unexpected findings in 10-week high-fat diet-fed mice, further efforts were made to demonstrate the relation between effector cell/transcription factor expression and their functional/cytokine production aspects in the mice after 15 weeks on a high-fat diet. Thus, expression of the Treg cell anti-inflammatory marker CD39, an ectonucleotidase and cleaves adenosine triphosphate (ATP), on Treg cells was decreased in plt-TGFβ$^{-/-}$ mice as compared to control mice (Supplementary figure 3).

#### ***Plt-TGFβ$^{-/-}$ elicits a pro-inflammatory profile in the aortic vessel wall***

Further efforts were made to reveal the impact of plt-TGFβ$^{-/-}$ on vascular inflammation in the aorta. Using RNA sequencing data from aortas of mice fed with a 15-week high-fat diet, gene ontology (GO) enrichment analyses showed that plt-TGFβ$^{-/-}$ enhanced lymphocyte and monocyte chemotaxis/infiltration and inflammatory responses (Supplementary figure 4A).

With proteomic data of the aorta from 15-week high-fat diet-fed mice, GO enrichment analyses demonstrated that plt-TGFβ$^{-/-}$ increased the expression of the major histocompatibility complex class II (MHC-II) protein complex (Supplementary figure 4B), indicating that the vascular inflammation was promoting local antigen presentation. KEGG pathway analyses revealed a strong enhancement of CD4 T effector cell responses, seen as enhanced Th1 and Th2 cell differentiation, Th17 cell differentiation, and T cell receptor signaling pathways. In agreement with the GO enrichment analyses, the KEGG analyses also indicated that plt-TGFβ$^{-/-}$ promoted antigen processing and presentation in the aorta, as well as NK cell-mediated cytotoxicity (Supplementary figure 4C).

### Supplementary Figures and Legends

#### Supplementary Figure 1

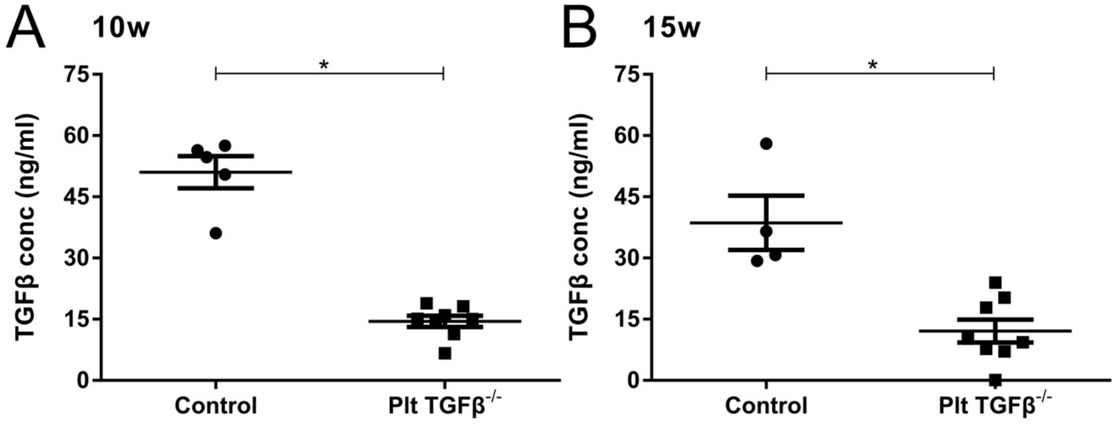


**Supplementary Figure 1. Platelet-specific TGFβ deficiency decreases plasma levels of TGFβ.** Plasma samples were prepared from blood collected from platelet-specific TGFβ1 deficient (Plt-TGFβ$^{-/-}$) mice and their littermate controls. The TGFβ levels of the mice on high-fat chow for 10 weeks (panel A) and 15 weeks (panel B) were measured using an ELISA assay. Student's *t*-test, *$p$<0.05, n=4-8.

### Supplementary Figure 2

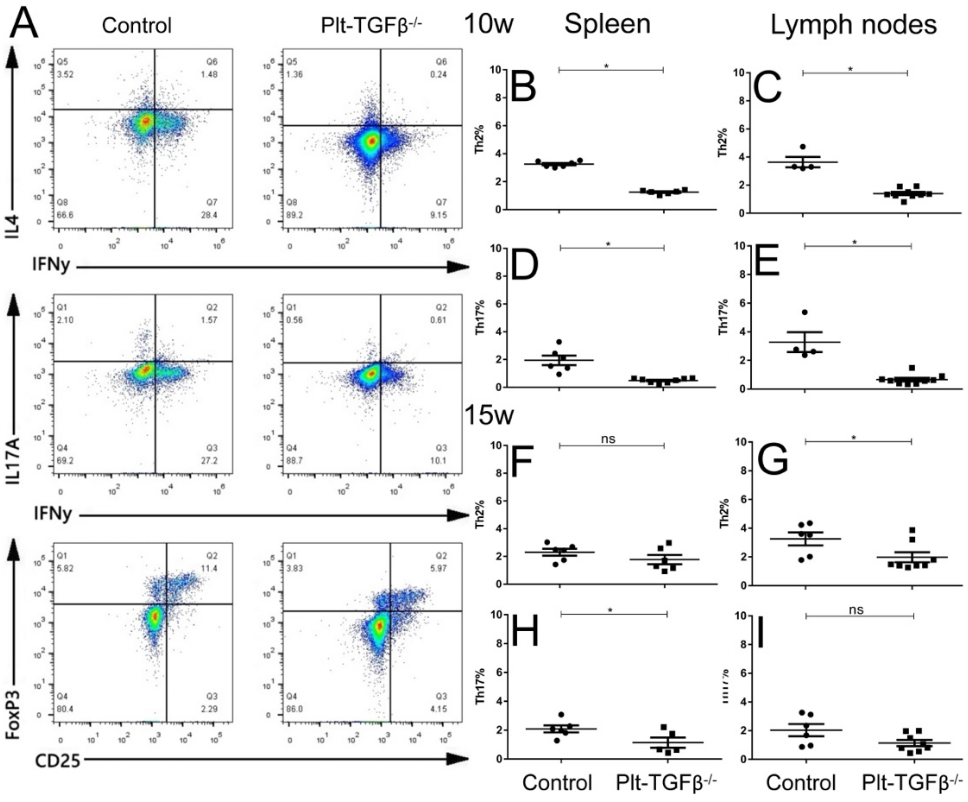


**Supplementary Figure 2. Platelet-specific TGFβ deficiency alters CD4$^+$ effector cell populations in the spleen and lymph nodes.** Plt-TGFβ$^{-/-}$ mice and their littermate control were fed with a high-fat chow for 10 and 15 weeks. The spleen and lymph nodes were harvested for the preparation of single cell suspensions at both time points. Panel A: Flow cytometric phenotyping of CD4$^+$ T effector cells, which were first stimulated using PMA (50 ng/ml) and ionomycin (5 µg/ml) for 2 hours, and followed by adding brefeldin A (5 µM) for further incubation for 4 hours. Representative pseudocolor plots. Panels B-C (10 W) and F-H (15 W): Th2 cells, CD4$^+$-IL-4$^+$. Panels D-E (10 W) and I-J (15 W): Th17 cells, CD4$^+$-IL-17A$^+$. *$p$<0.05 as compared to controls by Student's *t*-test, n=4-9.

**Supplementary Figure 3**

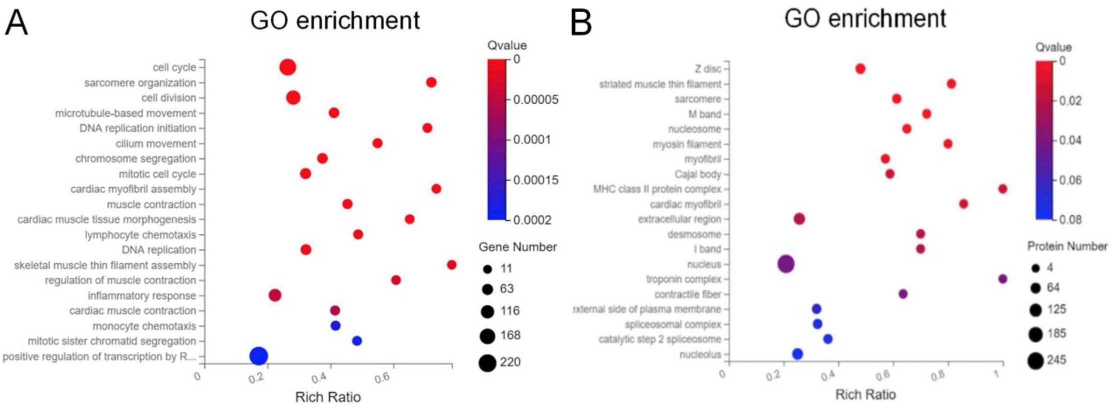


**Supplementary Figure 3. Altered transcriptomic and proteomic landscape in aortas from mice with platelet-specific TGFβ deficiency.**

Plt-TGFβ$^{-/-}$ mice and littermate controls were fed a high-fat diet for 15 weeks. The whole aorta was harvested, snap-frozen, and stored at -80°C for subsequent total RNA and protein isolation using Qiazol reagent and isopropanol precipitation. RNA sequencing was performed using the DNBSEQ™ Technology Platform, and mass spectrometric analyses and proteome profiling were carried out at BGI (Shenzheng, China). Data were analysed using the Dr. Tom program. Panel A: GO enrichment analysis of RNA sequencing. Panel B: GO enrichment analysis of aorta proteomic data. n=3-5.
